# Whole genome duplication enables rapid evolution of male-biased sex allocation in *Galax urceolata*

**DOI:** 10.1101/2025.06.19.660553

**Authors:** JH Williams, A Killeffer, C Bailey, B Cavalcante, T Birmley

## Abstract

Sex allocation in hermaphrodites is thought to evolve to a balance between fitness gained through male and female function. Whole genome duplication (WGD) might disrupt such gradually evolved patterns, since it has relatively instantaneous effects on sizes, but not numbers, of cells and organs. Here we ask whether sex allocation patterns differ between young neo-autotetraploid populations and their diploid progenitors within *Galax urceolata*.

- Floral organ sizes and numbers were measured using light microscopy and genome sizes verified with flow cytometry.
- Both cytotypes had the same number of flowers, anthers, and ovules per inflorescence, but floral organ sizes were proportionally longer in autotetraploids relative to diploid progenitors. Whole-plant allocation to volume of anthers increased by 176%, but of ovules only 70%. Autotetraploids produced 33% larger and 88% more pollen.
- WGD is known to double pollen volume, but pollen is biased to size reduction in young natural autotetraploids. In WGD-enlarged anthers, pollen size reduction allows increased pollen number to evolve without changing anther size. A literature review shows that higher pollen production in autopolyploid species is common, but not inevitable. Thus, we conclude that polyploidy can provide a pathway to increased pollen production, which may enhance male fitness during their early evolution.

## Introduction

According to sex allocation theory, hermaphroditic plants should invest separately in male and female functions until an optimal balance between the relative fitness gains through each sex is reached (Lloyd 1979; Charnov 1982; Burd 2025). Polyploidy, a recurrent and rapid evolutionary process, might disrupt such gradually evolved allocation patterns. Müntzing (1936) famously summarized a still prevailing view, that “the autotetraploid is mainly a magnified edition of the diploid,” based on a review of synthetic and natural intraspecific autotetraploids. In other words, the primary phenotypic effects of genome duplication are *gigas* effects - larger organs with conserved scaling relationships (allometry), and not much change in the numbers of parts. Such rapid organ size increases without decreases in organ numbers must incur substantial extra costs (Guignard et al. 2016; Corneillie et al. 2019), which presents the opportunity for differential partitioning of resources to evolve. Polyploidy may not directly alter sexual allocation to reproductive organs and gametes, but we know little about how and how fast differential allocation evolves in early stages of polyploid establishment.

Whole-genome duplication (WGD) occurs at the cell level, when non-reduction during mitosis or meiosis results in doubled chromosome numbers within a cell that is twice as large in volume as expected (Doyle and Coate 2019; Williams, unpublished). The roughly doubled volume of the meristematic cell in which WGD occurred is stably transmitted to its direct cellular descendants (Sax and Sax 1937; Satina et al. 1940; Bannan 1948). Perhaps the best-known effect of WGD in flowering plants is on pollen, a spore within which a haploid male gametophyte develops. “Pollen” of angiosperms are transiently free-living organisms that produce two sperm within a single “vegetative cell” during their short lifespan. In surveys of early generation synthetic autotetraploids, the vegetative cell of diploid (2n) pollen was consistently double the volume of 1n pollen of their diploid progenitors (Butterfass 1987; Altman et al. 1994; Westermann et al. 2024; Williams, unpublished). There are fewer studies on female gametophytes, but egg and central cell sizes are also roughly doubled in volume (Song et al. 2020).

Polyploidy, in contrast to WGD, is an organismal level phenomenon – each cell in a multicellular polyploid is far removed from its initial unreduced progenitor cell. In addition, most plant polyploids originate sexually by fusion of two gametes, at least one of which is unreduced (Harlan and de Wet 1975; De Storme and Geleen 2013). Since large genetic differences between progenitor gametes can generate heterosis (Birchler et al. 2010), the “purest” effects of WGD (due to doubled gene copy number and bulk DNA amount alone) are best seen in autopolyploids that have originated from closely related gametes.

The functional consequences of initially doubled cell size after WGD are immediate, so even the first generation of a multicellular autopolyploid has already experienced many cell generations of developmental selection (Buchholz 1922). A young polyploid might already display differentially altered developmental timing or duration of the cell cycles that give rise to different sexual organs and gametes. As evidence, the magnitude of cell enlargement seen in multicellular tissues of early generation, synthetic autotetraploids varies by cell type, genetic background, and species (Southworth and Pfahler 1992; Tsukaya 2013; del Pozo and Ramirez-Parra 2015; Robinson et al. 2018; Song et al. 2020).

Rapid compensation for *gigas* effects might occur as part of neo-polyploid stabilization, as exhibited in leaves and sepals of synthetic autotetraploids that are comprised of larger but fewer cells than their diploid progenitors (Robinson et al. 2018). Anther and ovary (carpel) sizes are also developmentally linked to the number and sizes of pollen grains and ovules, respectively. Pollen of neo-autotetraploids is genetically variable, since it can retain parental heterozygosity, whereas haploid pollen cannot (Williams 2021). Neo-autotetraploids are also likely to have reduced pollen fertility (Ramsey and Schemske 2002), resulting from large-scale mis-segregation of chromosome arms during male meiosis (Bomblies 2023). Thus, stabilization also involves intense selection for fertility restoration as well as on pollen performance during the first few generations after WGD, a period of maximal phenotypic and genetic variation of male gametophytes (Westermann et al. 2024). Less is known about the phenotypic effects of fertility restoration in female gametophytes, which develop heterotrophically within ovules.

Differential allocation to gamete sizes and numbers might evolve in response to the size effects of WGD, but to what degree and how rapidly? Sex allocation theory was founded on the premise that there should be trade-offs in allocation to male and female function when resources are limiting (Cruden 1977; Charnov 1982). Neo-polyploids must garner extra resources to support their larger biomass (Corneillie et al. 2019), and they are inevitably mate-limited (Stebbins 1971). Shifts in pollination efficiency or mating system can cause changes in the relative allocation of resources to pollen and ovules if there are fitness consequences (Cruden 1977; Harder and Johnson 2023). Successful establishment of neo-polyploids is more likely if their diploid progenitors were self-compatible and/or if they shifted to more inbred mating systems (Stebbins 1971; Barringer 2007; Husband et al. 2008). Shifts to more inbred mating systems are generally predicted to reduce allocation to male relative to female function (Cruden 1977; Lloyd 1987; Sicard and Lenhard 2011; Tsuchimatsu and Fujii 2022), usually with greater changes in pollen than in ovule number (Cruden 1977; Burd et al. 2009; Cunha et al. 2022; Cunha and Aizen 2023). Conversely, *gigas* effects in neo-polyploids involve enlargement of flowers and/or floral displays which can attract more pollinators (Goodwillie et al. 2010), facilitating increased outcrossing as well as greater gains to male over female fitness (Paterno et al. 2020; Cunha and Aizen 2023). Though ovule number generally evolves more slowly than pollen number (Lord 1980; Endress 2011; Cunha et al. 2022), it may increase as a function of floral (carpel) size (Yu et al. 2022) or by selection as variance in pollination efficiency increases (Burd 1995; Burd et al. 2009; Rosenheim et al. 2016).

In this study, we sought to examine the earliest effects of successful autotetraploidy on the sizes and numbers of floral parts in what is thought to be a young diploid-autotetraploid system, *Galax urceolata* (hereafter *Galax*, since it is monospecific). *Galax* is a mixed ploidy species that has many diploid, single-cytotype populations interdigitated with autotetraploid, single-cytotype populations in a broad zone of overlap. *Galax* was once called one of the best examples of incipient autopolyploidy (Stebbins 1950, based on Baldwin 1941), since tetraploids differ from diploids only by size-related traits. *Galax* has no close relatives, nor have strong geographic separation of cytotypes or substantial differences in niches evolved (Johnson et al. 2003; Gaynor et al. 2018). Both also share putative pollinators and flower concurrently at similar elevations (Barringer and Galloway 2017). Finally, tetraploid individuals and populations have formed repeatedly in the zone of overlap where diploids, triploids and tetraploids share similar genetic variation (Servick et al. 2015).

Here we focus narrowly on the question of how pre-pollination sex-allocation patterns differ between conspecific progenitor and derivative populations. Comparing sizes and numbers of mature structures provides a simple and relevant proxy for inferring changes in the *relative* energetic costs of sex allocation. Based on the conventional view that the effects of WGD are primarily cell and organ enlargement with conserved scaling, we reasoned that young autotetraploids, such as in *Galax,* would not yet have had time to evolve new sex allocation patterns. If altered sex allocation was observed, we expected it would have mainly arisen by differential compensation for *gigas* effects – i.e. numbers or sizes of organs of one sex would be more reduced that those of the other. Our specific questions were: 1) Are there differences in floral organ sizes between 2x and 4x plants and are floral organs isometric to each other within and between ploidy levels? 2) Are there differences in within-flower allocation to primary sexual organs between 2x and 4x plants? 3) Are there differences in whole-plant allocation to primary sexual organs between 2x and 4x plants? Finally, 4) Are there size-number trade-offs for flowers, anthers, pollen and ovules within and between cytotypes?

## Methods

### Study species

*Galax urceolata* (Diapensiaceae) is a perennial of the forest floor, commonly co-occurring with *Rhododendron* and *Kalmia* shrubs in the southern Appalachian region from Virginia to northern Georgia. There are two common cytotypes, a diploid (2n = 2x = 12) and a tetraploid (2n = 4x = 24), and populations of one or the other cytotype are found in a zone of overlap in the Blue Ridge mountains, predominantly in North Carolina. The base chromosome number in *Galax* is almost certainly x = 6, since n = 6 is the only number reported in four other genera in the family and is also the lowest number among several outgroup families (Rice et al. 2015). Thus, we refer to cytotypes/populations as being 2x or 4x, and pollen as being 1x or 2x, respectively. Support for autopolyploidy comes from lack of morphological differences other than size (Baldwin 1941), and similar flavonoids, allozymes, and DNA microsatellites (Soltis et al. 1983; Servick et al. 2015). Low inbreeding coefficients in 2x populations, the partitioning of genetic variation, and hand-pollination experiments all suggest that both cytotypes have more or less outcrossed mating systems (Servick et al. 2015; Barringer and Galloway 2017).

The flower of *G. urceolata* (Figure 1E,F) has five free sepals, five nearly free petals, and a fused androecium comprised of five fertile stamens alternating with five staminodes (Palser 1963). In mature *G. urceolata* flowers, the androecium forms a tube with five relatively spheroidal anthers that at maturity meet to form an opening above the stigma providing access to nectar at the base of the ovary. The ovary is tri-carpellate, with ca. 27-84 ovules per carpel, with a short, hollow style and a three-lobed stigma. Pollen grains are tri-aperturate with copious pollenkitt (Xi and Tang 1990) and both cytotypes are insect pollinated (Barringer and Galloway 2017). Close positioning of anthers above the stigma (reverse herkogamy) could also facilitate autonomous self-pollination, since anthers begin opening before flower opening and which overlaps morphological stigma receptivity. Isolated plants most commonly produce no or only one inflorescence each flowering season, otherwise 2-6. Flowering is acropetal (Figure 1C,D), from mid-May to mid-June/early July.

**Figure 1.**
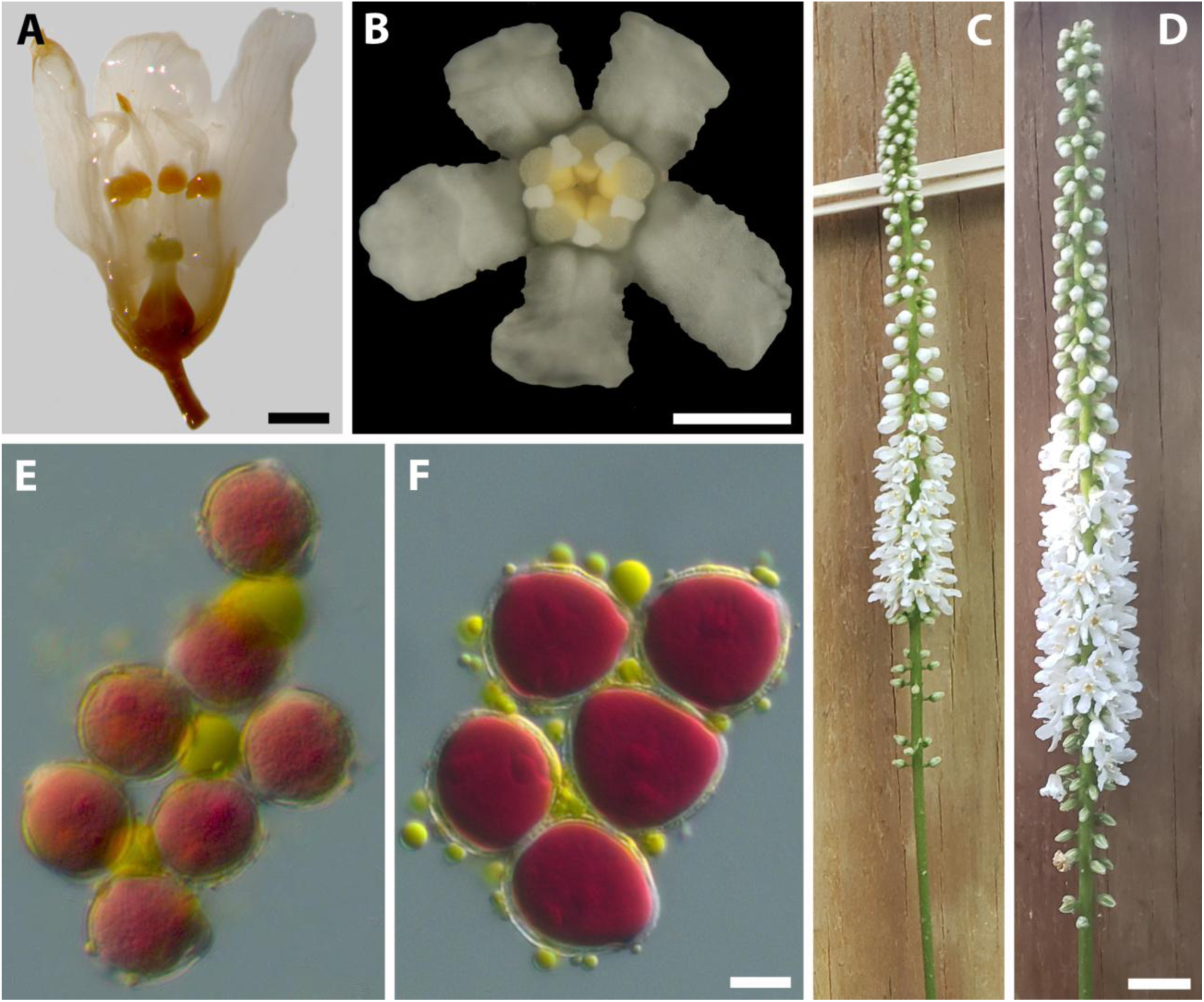
Relative sizes of *Galax urceolata* reproductive organs. A) Longitudinal view of flower at anthesis (2x, in Etoh); F) Face view of flower (2x, fresh). C, D) 2x and 4x inflorescences, respectively (same scale); E, F) 1x and 2x pollen, respectively (same scale); Scale bars: A = 1 mm; B = 0.5 mm; C,D = 1 cm; E, F = 10 *µ*m.

### Population sampling

Flowers and leaf tissue samples were collected from four wild populations of each ploidy in the Blue Ridge Mountains, USA (Table 1) in late May and early June, 2021 and 2022. Populations were near to or the same as previously identified single-cytotype populations, based on a flow cytometry survey by Servick et al. (2015). We later verified ploidy with flow cytometry (see results). Since tetraploid populations likely originated from nearby diploids, we sampled only within the southernmost (Blue Ridge mountain) of six diploid population genetic clusters identified by Servick et al. (2015).

**Table 1.** Study populations of *Galax urceolata.* All are in North Carolina, most in State forest, near BLR, Blue Ridge parkway. *N* is the number of plants sampled for floral traits or for ploidy. Putative cytotype was predicted from nearby single-cytotype populations assayed by Servick et al. (2015) (Servick site #).

| Pop | Lat/long | Elevation<br>(m) | Floral<br>*N* | Ploidy<br>*N* | Putative<br>cytotype | Servick<br>site # |
| --- | --- | --- | --- | --- | --- | --- |
| 6 | 35.57, -83.24 | 731 | 17 | 24 | 2x | 129 |
| 7 | 35.44, -83.11 | 1250 | 20 | 21 | 2x | 130 |
| 11 | 35.76, -81.92 | 340 | 21 | 04 | 2x | 125 |
| 12 | 35.76, -81.87 | 340 | 22 | 21 | 2x | Near 125 |
| 5 | 36.08, -81.83 | 1271 | 21 | 25 | 4x | Near 135 |
| 8 | 35.15, -82.82 | 1670 | 21 | 18 | 4x | 131 |
| 9 | 35.41, -82.75 | 1509 | 21 | 19 | 4x | 132 |
| 10 | 36.19, -81.74 | 1190 | 25 | 21 | 4x | Near<br>135, 136 |

Flowering individuals at least 3 meters apart were sampled along a transect. About half of the 165 plants sampled had only a single inflorescence (at least 47 2x and 36 4x genets). The other plants appeared to have more than one inflorescence, but it was not possible to determine if they came from the same genet. Inflorescences were photographed and flower numbers were counted from photos by counting the number of visible flowers plus the estimated number of hidden flowers. Estimated flower counts from photos were strongly correlated with actual flower counts on the same inflorescences (*r =* 0.97; Slope ± *SE =* 1.34 ± 0.10; *N* = 14 inflorescences).

### Measurement of floral organ sizes

Flowers of different ages were collected from inflorescences (Figure 1) and fixed in FAA for 24 h and then stored in 70% EtOH. Two floral ages were targeted: immature (just before opening) and mature (open with petals fully reflexed with bright yellow anthers and/or wet, whitish-green stigma). The flowers used in this study came from inflorescences at similar stages of phenology (Figure 1), scored as the proportion of flowers on an inflorescence that had progressed to or beyond the open flower stage. Mean (± *SE*) proportion of opened flowers on an inflorescence was: 2x = 0.57 ± 0.09 and 4x = 0.49 ± 0.09 (*P* = 0.56; *N* = 72, 82, respectively).

Mature flowers were cut longitudinally to expose the ovary and photographed in longitudinal view under the stereomicroscope at 6x magnification (Figure 1A). Pedicel width was measured just below the bulge of the outer surface of the receptacle. Curvilinear lengths of major floral organs were measured from apex to base where they attached to the receptacle (sepals, petals, stamens, pistils). Style length was measured from the roof of the upper ovarian cavity to stigma tip. Anthers are nearly spherical, so diameter (*D*) was measured from filament attachment point to tip, and then used to calculate volume as: *V =* π × *D^3^*/6. Anther-stigma distance was measured from the rear stamen, as the vertical distance from the bottom of anther to top of stigma (Figure 1A).

The same mature flowers were used to measure pollen (Figure 1E, F) and ovule sizes. Pollen grains were stained with acetocarmine and viability was assessed from 300 randomly chosen pollen grains (mean *N* = 305, 308 1x and 2x pollen grains, respectively) as: 1 minus the number of non-viable pollen per total pollen. Non-viable pollen lacked stain and/or was misshapen. Thirty viable pollen grains were randomly chosen and the minimum and maximum inner diameters (*D*) of the nearly spherical pollen (eg. vegetative cell size) were measured and averaged. Mean pollen size per flower was recalculated after removing putative 2n pollen, which were characterized as having a diameter > 1.25 times the mean diameter of pollen from that flower (a 1.26-fold increase in pollen diameter corresponds to a doubled volume of a sphere).

Pollen volume was calculated as: *V* = π x *D^3^*/6. Ovule area was directly measured by tracing outlines of three longitudinally oriented ovules from photos. Ovule volume was calculated as the volume of a sphere of diameter *D,* using the formula: *V* = π x *D^3^*/6, where *D* is the diameter of a circle of area, *A* = area of the measured ovule = π x (*D*/2)^2^.

Pollen and ovule numbers were counted on immature (nearly open) flowers, since mature flowers might have dispersed some pollen and/or aborted some ovules. Ovules were counted by dissecting one of three carpels and multiplying ovule number by three. Pollen number was estimated by crushing one anther per flower in a solution of 75 *µ*L acetocarmine with 10% glycerin and counting pollen grains on two replicate volumes of 0.1 *µ*L using a Hausser Scientific™ hemocytometer. Pollen number per flower was the average of the two replicates multiplied by 750 and by the number of anthers (usually five). Pollen viability was also determined from these counts as above (mean *N* = 23, 40 for 1x and 2x pollen grains, respectively). Total pollen viability was then calculated as the weighted average of pollen viabilities from both analyses. Pollen:ovule ratios were calculated for both total and viable-only pollen numbers per flower divided by total ovule number per flower. Whole plant allocation to volume was estimated for anthers, ovules, and pollen grains as: flower number per inflorescence x organ number per flower x organ volume. All photos were taken with a Zeiss Axiocam camera on a Zeiss Axioplan light microscope or stereoscope and image analyses were done using Zeiss Axiovision v. 8.2 software.

### Flow cytometry-

Relative ploidy level was inferred using flow cytometry (FCM) on fresh (< 2-day old) leaves. *Pisum sativum* cv. Ctirad was used as the standard, since the 2x *Galax* fluorescence peak was > 25% of the *P. sativum* G_1_/G_0_ peak and the 4x *Galax* G_1_/G_0_ peak did not overlap the *P. sativum* peak (based on formula in Temsch et al. 2022, p. 715). The standard peak fell between the G_1_/G_0_ and G_2_ peaks of 4x *Galax* (Supporting Information Table S1). Since collections were done in or near single-cytotype populations of Servick et al. (2015), we co-chopped ca. ¼ cm^2^ of fresh leaf material from each of 3 individuals with ca. ½ cm^2^ of the *Pisum* standard for each tube. If two sample G_1_/G_0_ peaks appeared, leaves were rerun individually. Samples were chopped in 1 mL of ice-cold Otto I buffer and filtered through a 35 *µ*m mesh. Tubes were centrifuged at 150g for 5 minutes, the supernatant removed, and 100 *µ*L of ice-cold Otto I buffer added. Tubes were incubated at room temperature for ∼ 30 minutes with occasional shaking before adding 0.8 mL of staining solution, prepared with 2 mL Propidium Iodide/Rnase (BD Pharmingen item # 550825), 20 mL PBS supplemented with 0.2% (w/v) bovine serum albumin, Ph 7.4 (BD Pharmingen stain buffer, item # 554657), and 4 *µ*L *β*-mercaptoethanol, filtered through a 0.2 *µ*m filter. Nuclei were incubated in the dark for 10-120 mins before running. FCM was run on a Cytek® Northern Lights flow cytometer (Cytek Biosciences, Fremont, CA), using the blue laser (excitation 488 nm at 50 mW) and optimal emission filter of 653-670 nm), to estimate 2C DNA content of leaf material. For each run, a minimum of 5000 events was observed, with a flow rate of 18-20 *µ*L min^-1^, gains of 100 (forward scatter) and 7 (side scatter), and a debris threshold of 200,000.

Data were analyzed using FCS Express 7 (De Novo Software 2024, Pasadena, CA). To estimate C-values in picograms (pg), we used the formula: Sample 2C-value = Standard 2C-value × (Sample G_1_/G_0_ fluorescence peak (RFUs)/Standard G_1_/G_0_ fluorescence peak (RFUs), where RFUs are relative fluorescence intensity units. Since the C-value of the standard has been revised several times, we report C-values based on three estimates of the true C-value of *Pisum sativum* cv. Ctirad: 9.09 pg (Doležel et al., 1998), 8.02 pg (Veselý et al., 2012), and 7.75 pg (Nix et al. 2024).

### Statistical analyses

After data collection, three 2x plants and two 4x plants were excluded from all statistical analyses, due to their exceptionally low pollen viabilities of 1%-6%. These could have been undetected odd-polyploids, aneuploids, or even newly formed tetraploids with meiotic problems, since pollen viability was high in all other plants (see results).

Each phenotypic trait was analyzed separately using a mixed model ANOVA, with Cytotype (2x vs. 4x) as a fixed effect and Population nested within Cytotype as a random effect (*N* = 4 populations for each cytotype). Because our primary goal was to estimate the effect of WGD on organ sizes within flowers, we controlled for variation in floral size, using log_10_ pedicel diameter (a proxy for local resource availability to flowers) as a covariate. Because pedicel width is correlated with ploidy level (see results), a violation of ANCOVA assumptions, we centered mean log_10_ pedicel diameters of each population on zero. Similarly, we included log_10_ flower number per inflorescence as a separate measure of whole-plant resource availability. Backward elimination of covariates and their interactions with cytotype was then used to drop non-significant parameters, with *P* ≥ 0.10 as the cut-off for elimination.

Use of pedicel size as a covariate also allowed us to test the allometric hypothesis that the relationship between floral organ sizes and flower (pedicel) size is linear and of similar slope in both cytotypes (Kilmer and Rodriguez 2016). A significant interaction effect would indicate that 2x and 4x cytotypes had different allometric slopes. For this reason, both organ length and pedicel width were log_10_-transformed for the analyses. All analyses used ordinary least squares and restricted maximum likelihood (REML) as implemented in JMP v. 16.0 (SAS Institute Inc. 2024). Effect sizes for each trait were calculated from the back-transformed LS means of each cytotype as: (4x LS mean)/(2x LS mean).

To explore the effect of ploidy level on organ-organ allometries a standard major axis (SMA) regression approach was used. In SMA regression, both variables are considered independent and log_10_ (organ A size) is linearly related to log_10_ (organ B size) as: log_10_(A) = *α* + *β* × log_10_(B), where *α* is the log_10_ intercept and *β* is the allometric slope (Warton et al. 2006). The orthogonal fit option in JMP with univariate variances was used, and 95% confidence intervals on the *β* of each cytotype were used to test the null hypothesis of isometry (*β* = 1).

Size-number trade-offs were examined with an ANCOVA approach to compare size-number relationships among cytotypes, testing the hypotheses of no trade-off (slope ≥ 0) and a perfect trade-off (slope = -1). Where warranted, the independent variable (eg. pollen number) was first adjusted for the effect of inflorescence or pedicel size (Vonhof and Harder 1995), if either had been significant as a covariate in a prior size or number analysis. Predicted values for each cytotype were calculated from the prior analysis, and then the mean for each cytotype was subtracted from each predicted value to estimate the effect of the covariate. In the trade-off analysis, the values of the independent variable were then adjusted for the covariate effect.

At the whole plant level, we tested for the effect of cytotype on total organ numbers and volumes per inflorescence, including pedicel diameter as a covariate. We also tested for intra-cytotype size-number trade-offs and differences in slopes among cytotypes, using a standard least squares approach to model organ size on organ number, with the full model including both measures of resource availability, log_10_ pedicel diameter and log_10_ flower number per inflorescence as above, and backward elimination of non-significant parameters (*P* > 0.10).

Finally, to test for differences in primary sexual allocation patterns between cytotypes at the whole plant level, we used a two-way ANOVA with Gender (male vs. female organs) and Cytotype (2x vs. 4x) as fixed effects. In the case of a significant interaction effect, we used pre-planned contrasts to test for differences between cytotypes within male organs (anthers or pollen grains) or female organs (ovules).

## Results

### Genome sizes and ploidy levels of cytotypes

*Galax urceolata* and the *Pisum sativum* var. Ctirad standard leaves were run separately and together, and when the standard was run with either one 2x, one 4x, or both a 2x and a 4x leaf, the 2C peaks of both samples and the standard (G_0_/G_1_ of the cell cycle) remained at very similar positions (Figure S1). The mean G_0_/G_1_ sample to standard ratios were 0.296 and 0.608, for 2x and 4x populations, respectively.

All 73 plants from putative 4x populations had similar relative fluorescence peaks with a coefficient of variation (CV) of < 2% and an intensity approximately double that of the 2x mean from 70 plants in putative 2x populations (CV’s < 4%). The mean 2n genome size (GS) of 2x populations was 2.292 ± 0.017 pg, whereas the mean 2n GS of 4x populations was 4.714 ± 0.015 pg (Table 2, based on Nix et al. (2024) estimate of *Pisum sativum* cv. Ctirad genome size). The 4x genome size was 2.8% larger than predicted based on doubled 2x genome size (Table 2). Importantly, the predicted genome size for triploids was outside of and intermediate to the upper and lower limits of the 2x and 4x ranges, respectively (Table 2).

**Table 2.** 2C genome size estimates of *Galax urceolata* diploid (2x) and tetraploid (4x) populations. For comparison, genome sizes were calculated using three alternative estimates of the 2C DNA mass of the standard, *Pisum sativum* cv Ctirad.

| Population | N plants | Sample/<br>standard<br>ratio | Sample/<br>standard<br>CV (%) | Nix et al. 2024 |  | Veselý et al., 2012 |  | Doležel et al., 1998 |  |
| --- | --- | --- | --- | --- | --- | --- | --- | --- | --- |
|  |  |  |  | Mean ± SD<br>(pg) | Min/max<br>(pg) | Mean ± SD<br>(pg) | Min/max<br>(pg) | Mean ± SD<br>(pg) | Min/max<br>(pg) |
| 6 | 24 | 0.298 | 3.72 | 2.309 ± 0.086 | 1.995/2.624 | 2.391 ± 0.089 | 2.066/2.718 | 2.709 ± 0.101 | 2.341/3.079 |
| 7 | 21 | 0.295 | 3.63 | 2.286 ± 0.083 | 2.152/2.441 | 2.368 ± 0.086 | 2.229/2.529 | 2.683 ± 0.097 | 2.525/2.865 |
| 11 | 4 | 0.295 | 1.99 | 2.286 ± 0.045 | 2.225/2.256 | 2.368 ± 0.047 | 2.305/2.337 | 2.683 ± 0.053 | 2.611/2.647 |
| 12 | 21 | 0.295 | 3.63 | 2.285 ± 0.083 | 1.995/2.326 | 2.367 ± 0.086 | 2.066/2.409 | 2.682 ± 0.097 | 2.341/2.730 |
| Mean 2x | 70 | 0.296 |  | 2.292 ± 0.017 |  | 2.374 ± 0.018 |  | 2.689 ± 0.02 |  |
| 5 | 23 | 0.609 | 1.75 | 4.716 ± 0.082 | 4.591/4.984 | 4.885 ± 0.085 | 4.756/5.163 | 5.535 ± 0.097 | 5.388/5.849 |
| 8 | 21 | 0.607 | 1.71 | 4.703 ± 0.080 | 4.613/4.795 | 4.871 ± 0.083 | 4.778/4.967 | 5.519 ± 0.094 | 5.413/5.627 |
| 9 | 19 | 0.609 | 1.66 | 4.719 ± 0.078 | 4.584/4.825 | 4.889 ± 0.081 | 4.749/4.998 | 5.538 ± 0.092 | 5.380/5.662 |
| 10 | 22 | 0.609 | 1.73 | 4.719 ± 0.081 | 4.459/4.984 | 4.888 ± 0.084 | 4.618/5.163 | 5.538 ± 0.096 | 5.232/5.849 |
| Mean 4x | 85 | 0.608 |  | 4.714 ± 0.015 |  | 4.883 ± 0.016 |  | 5.532 ± 0.018 |  |
| 3x exp. value<sup>a</sup> |  |  |  | 3.437 |  | 3.561 |  | 4.034 |  |
| 4x exp. value |  |  |  | 4.583 |  | 4.747 |  | 5.378 |  |

<sup>a</sup> Note that the expected 3C-values (1.5 x 2C mean) are outside the 2C and the 4C ranges.

### The effect of ploidy on floral organ sizes-

Ploidy level had a significant positive effect on lengths of all major floral organs, after controlling for flower size (pedicel diameter) and/or inflorescence size (flower number) (Table 3; Figure 1). Pedicel size had the strongest effect on lengths of petals, anthers, and styles, whereas flower number per inflorescence only appeared as a factor in anther length. 4x sepals, petals and pistils were 34-35% longer than in 2x plants, whereas 4x stamens were only 24% longer. Since separate slopes models were rejected in all cases (interaction effects ranged from *P* = 0.36-0.83), there was no evidence that the scaling of major floral organs relative to flower or inflorescence size differed among cytotypes. The 95% confidence intervals of the random effect in each model always included zero and variation in trait means among populations within cytotypes was minor relative to variation among cytotypes.

**Table 3.** Floral trait dimensions of 2x and 4x cytotypes of *G. urceolata.* Values are back-transformed least-squares (LS) means (bolded) centered within their 95% confidence intervals, and effect size is the ratio of 4x to 2x back-transformed LS means. *P* values are from a same slopes ANCOVA model for the main effect of cytotype. The log-log slopes of covariates *β_ped_* (pedicel diameter) and *β_flwr_* (flower number) are shown if included in model (*P* < 0.10). With minor exception, *N* = 76, 84 for 2x and 4x cytotypes, respectively. A-S Distance, anther-stigma distance; L, length; V, volume. All measurements are in mm.

| | Trait | 2x | 4x | Effect size | $R^2_{adj.}$ | $P_{cytotype}$ | Log-log $\beta^a$ |
| --- | --- | --- | --- | --- | --- | --- | --- |
| 1 | Pedicel D | {0.33, **0.35**, 0.37} | {0.49, **0.52**, 0.56} | **1.48** | **0.67** | **<0.0001** |  |
| 2 | Sepal L | {1.52, **1.67**, 1.83} | {2.05, **2.25**, 2.47} | **1.35** | **0.62** | **0.0017** |  |
| 3 | Petal L | {3.90, **4.07**, 4.24} | {5.22, **5.45**, 5.68} | **1.34** | **0.69** | **<0.0001** | $\beta_{ped} = +0.30****$ |
| 4a | Stamen L | {2.68, **2.91**, 3.16} | {3.32, **3.60**, 3.91} | **1.24** | **0.51** | **0.0044** | $\beta_{ped} = +0.20**$ |
| 4b | Filament L | {2.36, **2.57**, 2.81} | {2.86, **3.13**, 3.42} | **1.22** | **0.41** | **0.0090** | $\beta_{ped} = +0.19*$ |
| 4c | Anther L | {0.30, **0.33**, 0.37} | {0.42, **0.46**, 0.51} | **1.39** | **0.71** | **0.0009** | $\beta_{ped} = +0.27***$<br>$\beta_{flwr} = +0.11**$ |
| 5a | Pistil L | {1.44, **1.54**, 1.64} | {1.94, **2.08**, 2.22} | **1.35** | **0.63** | **0.0003** | $\beta_{ped} = +0.24***$ |
| 5b | Ovary L | {0.70, **0.80**, 0.91} | {0.90, **1.02**, 1.17} | **1.28** | **0.42** | **0.0175** |  |
| 5c | Style L | {0.68, **0.73**, 0.78} | {0.96, **1.03**, 1.10} | **1.41** | **0.56** | **0.0001** | $\beta_{ped} = +0.41****$ |
| 6 | A-S Distance | {0.35, **0.45**, 0.59} | {0.34, **0.44**, 0.58} | 0.97 | 0.15 | 0.8717 | $\beta_{ped} = +0.83**$ |
| 7 | Stam L-Pistil L | {1.22, **1.36**, 1.52} | {1.35, **1.51**, 1.68} | 1.11 | 0.20 | 0.1573 | $\beta_{ped} = +0.17$ |

<sup>a</sup> Significance levels: no asterisk, $P < 0.10$ ; \*, $P < 0.05$ ; \*\*, $P < 0.01$ ; \*\*\*, $P < 0.001$ , \*\*\*\* $P < 0.0001$ .

The scaling relationships among 2x floral organs were mostly isometric, and similar to the pattern among 4x organs (Table 4). In all 6 pairs of SMA organ-organ regressions, the 95% confidence intervals (*CI*_0.95_) on the allometric log-log slopes of 4x cytotypes broadly overlapped those of 2x cytotypes (Table 4). Furthermore, all but one *CI*_0.95_ on slopes included one, consistent with conservation of isometry across cytotypes. The *CI*_0.95_ of the pistil-petal regression for the 4x cytotype did not include 1, but it broadly overlapped the 2x slope estimate (Table 4).

**Table 4.**
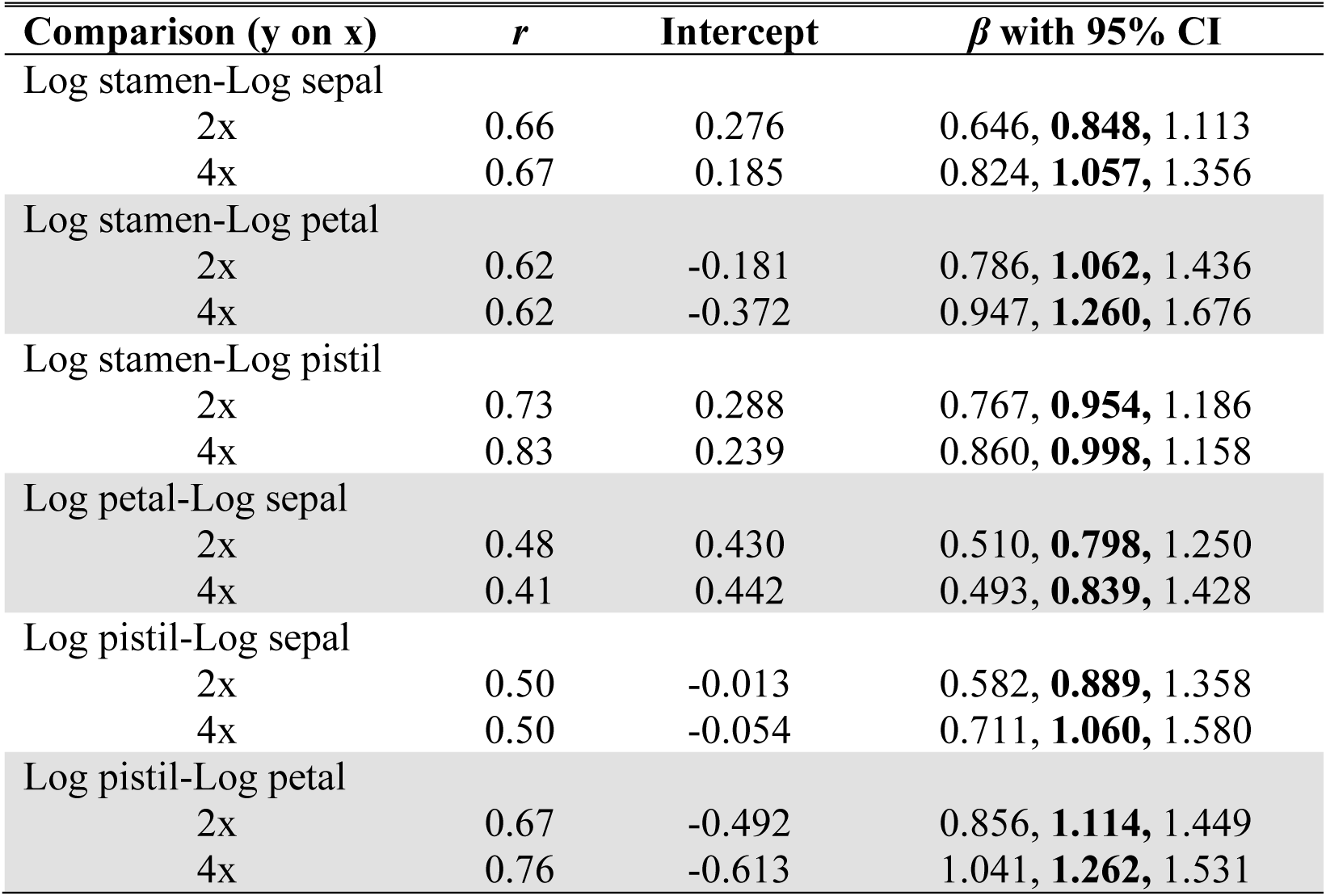
Standard major axis regressions on major floral organs in 2x and 4x *G. urceolata.* Isometry is indicated by an allometric slope of ***β*** =1. ***β*** > 1, indicates that organ size on the y-axis increased in size faster than organ size on the x-axis.

Although there was no evidence for differences in allometric slopes between cytotypes, when 2x and 4x plants were pooled as a single population, the three SMA regressions involving stamens had slopes significantly < 1 (Figure 2). This result reflects the fact that flower enlargement by WGD resulted in a smaller size increase of stamens than expected, even though the allometric slopes within cytotypes for these same comparisons were isometric and similar (Table 4). Sepals, petals, and pistils were isometric to each other (Figure 2). In sum, flower enlargement by WGD did not alter the proportions among floral organs other than stamens.

**Figure 2.**
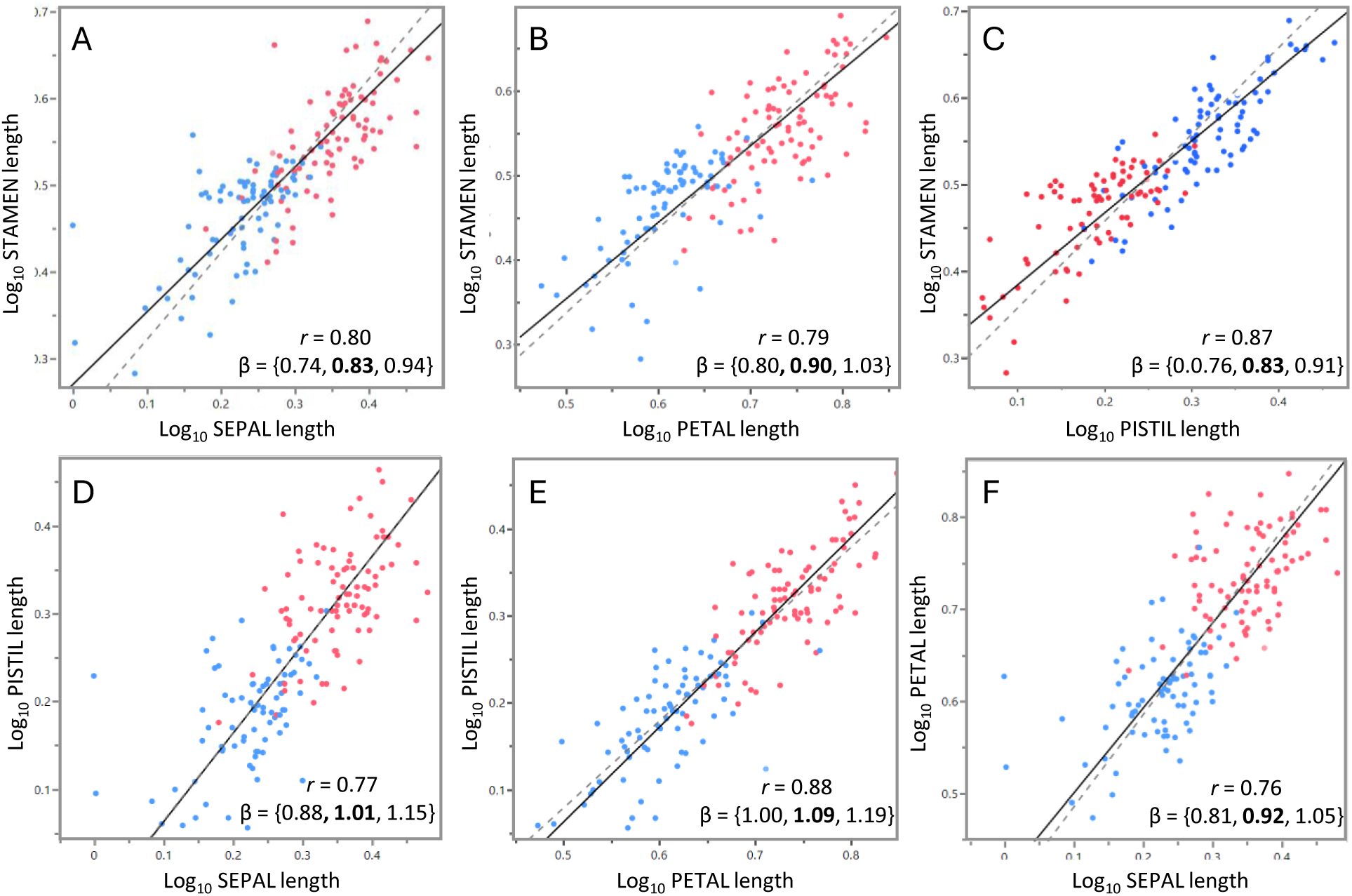
Allometry of floral organs in *Galax urceolata.* Each point is one flower on one plant *(N* = 160). Diploids, blue; autotetraploids, red. Solid line is the fitted SMA regression line, whereas dashed line represents isometry (slope of one, centered on the means). The slope, β, is bolded within its 95% confidence interval.

Autotetraploid stamens were proportionally shorter than other 4x floral parts because their filaments were less elongated – a 22% increase instead of the 34-35% seen in other major organs (Table 3). Importantly, the diameter of pollen-producing anthers was 39% greater in 4x than 2x plants, whereas the length of ovule-producing ovaries was only 28% greater (styles were 41% longer, such that 4x pistils had a comparable length increase to petals and sepals).

We expected anther-stigma distance to increase with floral size, however, neither anther-stigma distance (*P* = 0.87) nor the difference between stamen and pistil length (*P* = 0.16) were affected by ploidy level (Table 3). But floral size did increase anther-stigma distance within cytotypes, as indicated by a large positive pedicel diameter effect (Table 3). Thus, despite the more or less isometric size increases of 4x sepals, petals, pistils, anthers and styles, anther-stigma distance was conserved because 4x filaments were proportionally less elongated relative to all other floral organs.

### The effect of ploidy on primary sexual organs and gametophytes

An ovule is an integumented megasporangium that develops a single female gametophyte that produces a single egg. Ovule volumes were 62% larger in 4x than in 2x plants, but both cytotypes produced about 46 ovules per flower (Table 5). Anthers are microsporangia, which house male gametophytes (pollen grains), each of which produces two sperm. The number of anthers per flower was 5.0 versus 5.05 in 2x and 4x plants, respectively (*P* = 0.37). However, the 39% greater length of 4x over 2x anthers reflects a 167% increase in anther volume (outside diameter), indicating a much greater capacity to house pollen grains (Table 5).

**Table 5.**
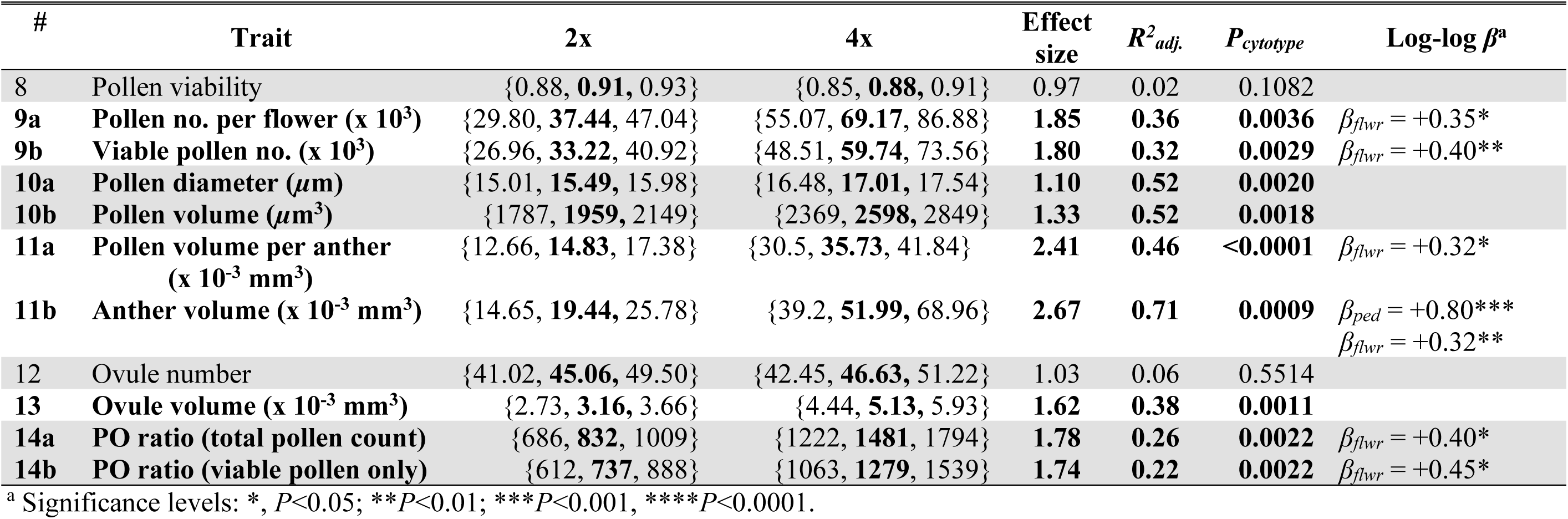
Gamete-bearing organ numbers and dimensions of 2x and 4x cytotypes of *G. urceolata.* Values are back-transformed least-squares (LS) means (bolded) centered within their 95% confidence intervals, and effect size is the ratio of 4x to 2x back-transformed LS means. *P* values are from a same slopes ANCOVA model for the main effect of cytotype. The log-log slopes of covariates *β_ped_* (pedicel diameter) and *β_flwr_* (flower number) are shown if included in model (*P* < 0.10). With minor exception, *N* = 76, 84 for 2x and 4x cytotypes, respectively. Pollen viability is the proportion of pollen scored as viable. PO ratio, pollen to ovule ratio.

| # | Trait | 2x | 4x | Effect size | $R^2_{adj.}$ | $P_{cytotype}$ | Log-log $\beta^a$ |
| --- | --- | --- | --- | --- | --- | --- | --- |
| 8 | Pollen viability | {0.88, **0.91**, 0.93} | {0.85, **0.88**, 0.91} | 0.97 | 0.02 | 0.1082 |  |
| 9a | Pollen no. per flower (x 10<sup>3</sup>) | {29.80, **37.44**, 47.04} | {55.07, **69.17**, 86.88} | **1.85** | **0.36** | **0.0036** | $\beta_{flwr} = +0.35^*$ |
| 9b | Viable pollen no. (x 10<sup>3</sup>) | {26.96, **33.22**, 40.92} | {48.51, **59.74**, 73.56} | **1.80** | **0.32** | **0.0029** | $\beta_{flwr} = +0.40^{**}$ |
| 10a | Pollen diameter ($\mu m$) | {15.01, **15.49**, 15.98} | {16.48, **17.01**, 17.54} | **1.10** | **0.52** | **0.0020** | |
| 10b | Pollen volume ($\mu m^3$) | {1787, **1959**, 2149} | {2369, **2598**, 2849} | **1.33** | **0.52** | **0.0018** | |
| 11a | Pollen volume per anther (x 10<sup>-3</sup> mm<sup>3</sup>) | {12.66, **14.83**, 17.38} | {30.5, **35.73**, 41.84} | **2.41** | **0.46** | **<0.0001** | $\beta_{flwr} = +0.32^*$ |
| 11b | Anther volume (x 10<sup>-3</sup> mm<sup>3</sup>) | {14.65, **19.44**, 25.78} | {39.2, **51.99**, 68.96} | **2.67** | **0.71** | **0.0009** | $\beta_{ped} = +0.80^{***}$<br>$\beta_{flwr} = +0.32^{**}$ |
| 12 | Ovule number | {41.02, **45.06**, 49.50} | {42.45, **46.63**, 51.22} | 1.03 | 0.06 | 0.5514 |  |
| 13 | Ovule volume (x 10<sup>-3</sup> mm<sup>3</sup>) | {2.73, **3.16**, 3.66} | {4.44, **5.13**, 5.93} | **1.62** | **0.38** | **0.0011** |  |
| 14a | PO ratio (total pollen count) | {686, **832**, 1009} | {1222, **1481**, 1794} | **1.78** | **0.26** | **0.0022** | $\beta_{flwr} = +0.40^*$ |
| 14b | PO ratio (viable pollen only) | {612, **737**, 888} | {1063, **1279**, 1539} | **1.74** | **0.22** | **0.0022** | $\beta_{flwr} = +0.45^*$ |

<sup>a</sup> Significance levels: \*, $P < 0.05$ ; \*\*, $P < 0.01$ ; \*\*\*, $P < 0.001$ ; \*\*\*\*, $P < 0.0001$ .

Autotetraploids produced 85% more pollen per flower (or 80% more viable pollen) than diploids, and such 2x pollen grains were 33% larger in volume than the 1x pollen of their diploid relatives (Table 5; Figure 1E, F). The viability of 2x pollen was not statistically different from that of 1x pollen, consistently near 90%. The total volume of pollen protoplasm (inside volume) produced per 4x anther was more than double (141%) of that produced per 2x anther, comparable to the more-than-doubled volume change recorded for anthers (Table 5).

In terms of within-flower allocation to gamete numbers, the mean pollen-ovule (PO) ratio increased by 78%, from 832 in 2x flowers to 1,481 in 4x flowers (Table 5; 74% higher for viable pollen). Since 2x and 4x flowers had similar ovule and anther numbers, the higher PO ratios of 4x plants were solely a consequence of greater pollen number per anther.

### The effect of ploidy on whole plant allocation of primary sexual traits

Sixty-three percent of 2x and 58% of 4x plants in our sample were isolated plants that produced only a single inflorescence. For the remainder, inflorescence numbers could not be determined accurately, since we could not distinguish ramets from genets. Statistics on the single-inflorescence dataset were almost identical to those reported below for the full dataset (Supporting Information Tables S1, S2).

The number of flowers per 2x and 4x inflorescences was similar (107 vs. 105, respectively; *P* = 0.58; Table 6). The total volume of ovules per inflorescence was 70% greater in 4x than in 2x plants, whereas the total volume of anthers increased by 176% (Table 6). Since numbers of anthers (microsporangia) and ovules (megasporangia) per flower were not affected by ploidy level (Tables 3, 4), the greater total allocation to both anthers and ovules at the whole-plant level was due solely to size effects. Importantly, the proportional increase in total anther volume of 4x plants was significantly greater than the increase in ovule volume, as indicated by a significant Cytotype x Gender interaction (two-way ANOVA, *P* < 0.0001; Figure 3A).

**Table 6.** Whole-plant sexual allocation patterns of 2x and 4x cytotypes of *G. urceolata.* Values are back-transformed least-squares (LS) means (bolded) centered within their 95% confidence intervals, and effect size is the ratio of 4x to 2x back-transformed LS means. For each trait, *P* values are from a same slopes ANCOVA model for the main effect of cytotype. The log-log slope of the covariate, pedicel diameter (D) is shown if included in model (*P* < 0.10). With minor exception, *N* = 76, 84 for 2x and 4x cytotypes, respectively.

| # | Trait (per inflorescence) | 2x | 4x | Effect size | $R^2_{adj.}$ | $P_{cytotype}$ | Log-log $\beta^a$ |
| --- | --- | --- | --- | --- | --- | --- | --- |
| 15 | Total flower number | {98.4, **104.8**, 111.7} | {100.6, **107.1**, 114.0} | 1.02 | -0.03 | 0.5801 | NA |
| 16 | Total anther volume (mm<sup>3</sup>) | {7.37, **10.17**, 14.03} | {20.34, **28.06**, 38.70} | **2.76** | **0.57** | **0.0016** | $\beta_{ped} = +2.26^{**}$ |
| 17a | Total pollen number (x 10<sup>3</sup>) | {3074, **3930**, 5025} | {5795, **7408**, 9469} | **1.88** | **0.27** | **0.0045** |  |
| 17b | Viable pollen (x 10<sup>3</sup>) | {2775, **3478**, 43603} | {5098, **6387**, 8001} | **1.84** | **0.22** | **0.0037** |  |
| 18a | Total pollen volume (mm<sup>3</sup>) | {6.44, **7.80**, 9.45} | {15.89, **19.24**, 23.31} | **2.47** | **0.37** | **0.0002** | $\beta_{ped} = +0.58$ |
| 18b | Viable pollen vol. (mm<sup>3</sup>) | {5.70, **6.87**, 8.29} | {13.80, **16.64**, 20.05} | **2.42** | **0.33** | **0.0002** |  |
| 19 | Total ovule number | {4313, **4774**, 5285} | {4499, **4992**, 5539} | 1.05 | <0.01 | 0.4494 |  |
| 20 | Total ovule volume (mm<sup>3</sup>) | {12.29, **14.99**, 18.29} | {20.92, **25.51**, 31.09} | **1.70** | **0.29** | **0.0036** |  |

<sup>a</sup> Significance levels: no asterisk, $P < 0.10$ ; \*, $P < 0.05$ ; \*\* $P < 0.01$ ; \*\*\* $P < 0.001$ , \*\*\*\* $P < 0.0001$ .

**Figure 3.**
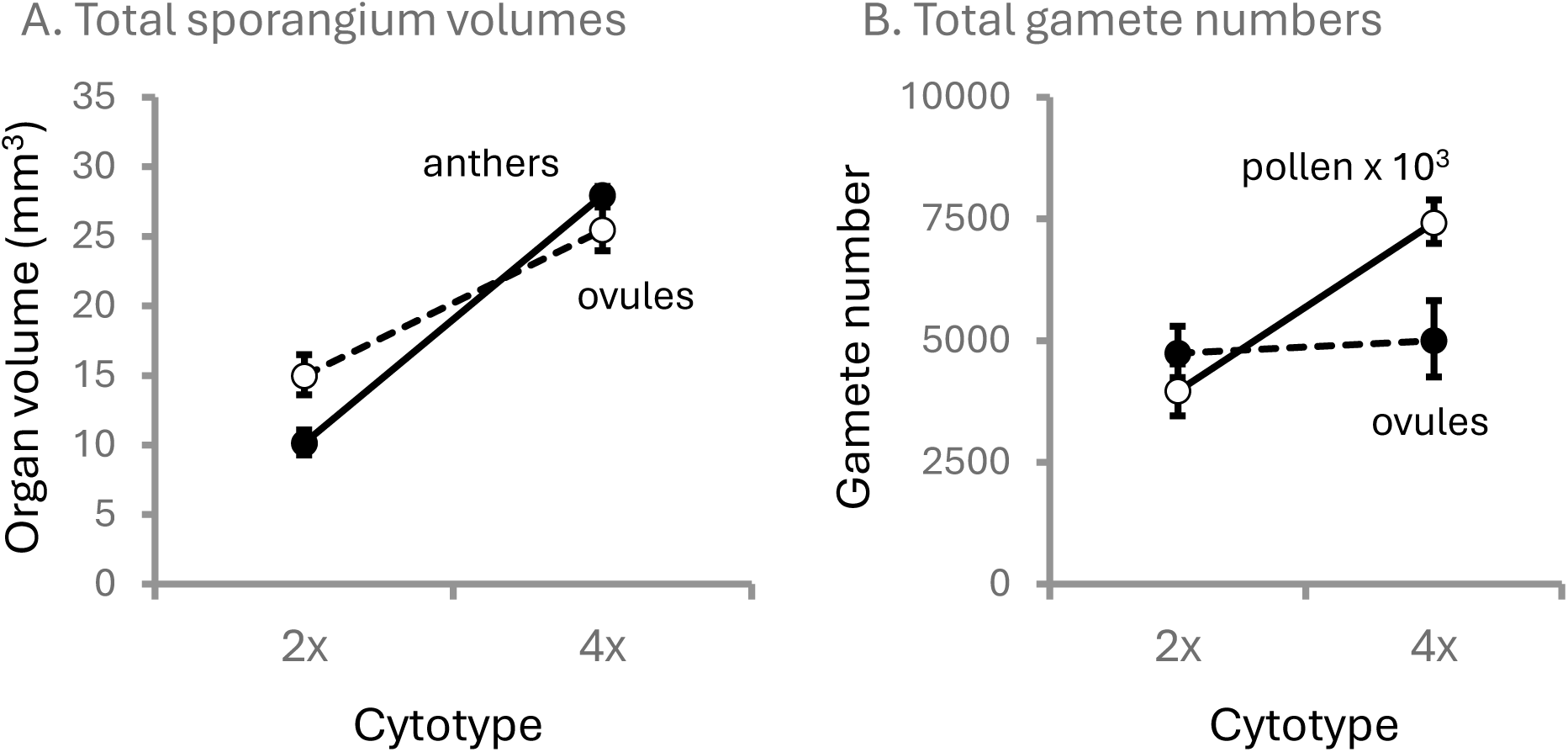
Whole-plant sexual allocation in 2x and 4x cytotypes of *Galax urceolata.* A) Total volumes of gamete-bearing organs per inflorescence. B) Total gamete numbers per inflorescence. Pollen and ovule numbers are proxies for gamete numbers, since each contributes one gamete to the next generation. Means and error bars (95% confidence limits) are back-transformed values. Pollen numbers are x10^3^ for graphical purposes

Autopolyploids shifted more resources to male function in terms of volumes, but they also increase pollen production far more than ovule production. At the whole-plant level, autotetraploids produced 88% more pollen grains, but only 5% more ovules, than 2x inflorescences (two-way ANOVA interaction effect, *P* < 0.0001; Figure 3B). The statistical results were the same when viable, instead of total, pollen numbers were used (4x inflorescences produced 84% more viable pollen). Thus, within sexes, there was no evidence for compensation of increased size by reduced numbers - neither size nor number of anthers, pollen or ovules decreased with ploidy. But there was an among cytotype trade-off in sexual function – total anther volume increased proportionally more than total ovule volume.

For trade-offs within cytotypes, the best model testing pollen size on pollen number, after dropping the interaction effect (*P* = 0.77), found a common log-log pollen size-number slope of - 0.385 (Figure 4). The 95% confidence interval on the slope slightly overlapped zero, but not -1 (95% CI: -0.850, 0.081). For ovule size on number, all effects were non-significant (*R^2^_adj._* = 0.01), with the strongest effect being the interaction (*P* = 0.056). The log-log slopes were +0.062 (*P* = 0.397) for 2x and -0.164 (*P* = 0.059) for 4x cytotypes. For flower size (pedicel diameter) on flower number per inflorescence, all effects were non-significant (*R^2^_adj._* = 0.0; log-log slope = 0.186; *P* = 0.204).

**Figure 4.**
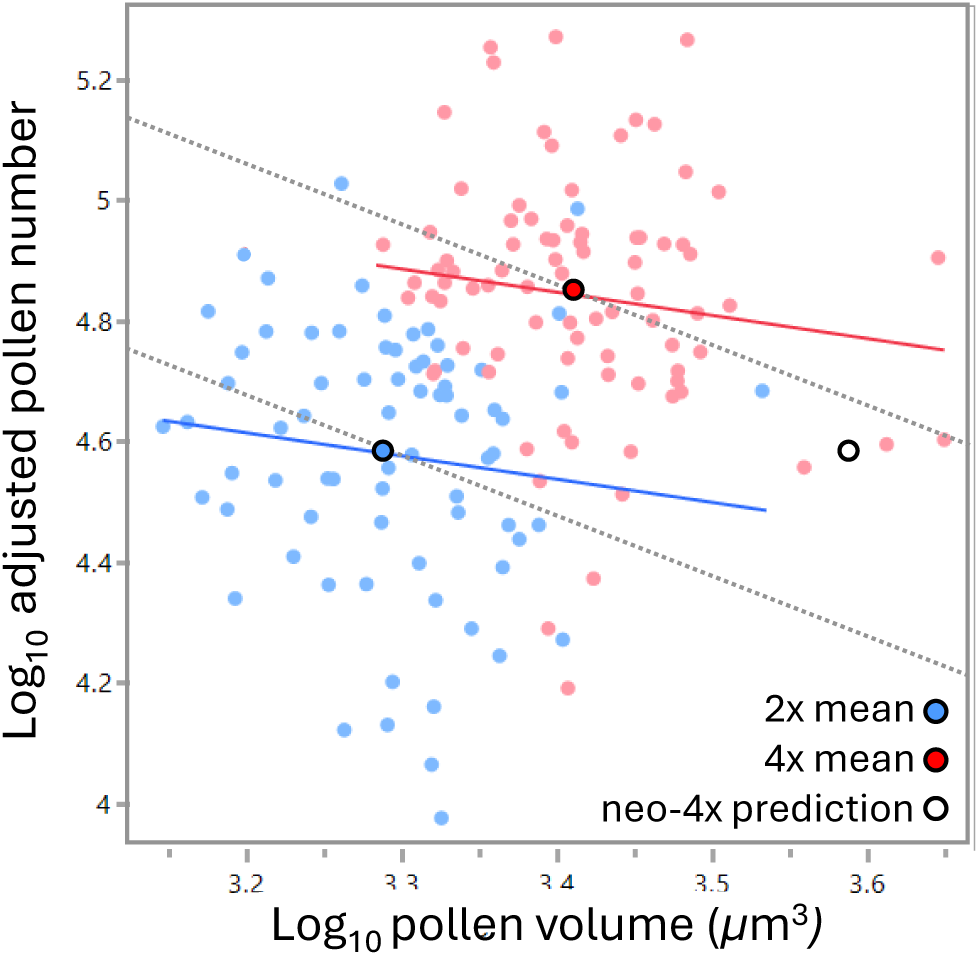
Pollen size-number relationships in 2x and 4x cytotypes of *Galax urceolata*. Blue, 2x flowers; Red, 4x flowers. Log pollen number was adjusted after accounting for the effect of inflorescence size on pollen number (see methods). At inception, neo-4x autopolyploids are expected to have doubled pollen grain volume without change in pollen number (unfilled circle). Dashed lines represent isoclines with slope of -1 (any point along each dashed line has the same total volume of pollen per flower as the 2x or 4x means).

## Discussion

The primary phenotypic outcome of whole genome duplication is cell enlargement, whereas multicellular polyploids experience downstream effects of WGD on the development of organs and gametes. Because allocation to sizes and numbers of sexual organs in bisexual flowers involves semi-independent developmental pathways and construction costs, we hypothesized that sex allocation patterns might be altered during polyploid stabilization and establishment.

Intraspecific autotetraploids in *Galax* had doubled 2x genome size, large floral organ size effects, conserved scaling of floral organs, and no evidence for changes in numbers of flowers, major floral organs, anthers, or ovules - all indicators of the predominant effects of WGD, and not of substantial post-WGD evolution. On the other hand, whole-plant investment in volumes of primary sexual organs of autotetraploids relative to diploid progenitors, increased by 176% for anthers, but only by 70% for ovules. The greater increase in 4x anther volume compared to ovule volume was reflected in a dramatic 88% increase in pollen (sperm) number with no corresponding change in ovule (egg) number. We attribute this altered sexual allocation pattern to post-WGD processes that have acted rapidly to differentially modify the primary size effects of WGD on male and female function, as discussed below.

### Polyploidy affected floral organ sizes, not numbers

Polyploidy was originally recognized when trying to explain the rare occurrence of plants with excessively large buds and flowers (Ramsey and Ramsey 2014). Floral *gigas* phenotypes were later reported on both newly synthesized and natural intraspecific autotetraploids (Müntzing 1936). The flowers of 4x *Galax* plants were larger by every measure of flower size -pedicel diameters, and sepal and petal lengths. Furthermore, 4x floral organ lengths all increased by a similar magnitude, such that organ-organ scaling in 2x flowers was largely retained. Conserved scaling of parts suggests that genetic correlations among organs were not much affected by WGD. Furthermore, the magnitude of size changes of major organs, 34-35%, is similar or greater than the magnitude of size changes seen at inception in synthetic neo-autotetraploids (Porturas et al. 2019; Schmickl et al. 2024).

On the other hand, flower number per inflorescence and organ numbers per flower were not altered by polyploidy. These results are consistent with studies of meristic reproductive traits that show synthetic autotetraploids to have similar or fewer numbers of inflorescences (8 studies) or flowers/florets (16 studies) than their diploid progenitors (Porturas et al. 2019). Major floral organs arise from a determinate floral apical meristem (FM) and in most monocots and eudicots floral part numbers are strongly canalized (Endress 2011). Canalization is supported by the fact that meristem enlargement by WGD did not change the numbers of sepals, petals, stamens or carpels. The phyllotactic pattern at the FM constrains anther number but not ovule number.

Ovule primordia are initiated long after formation of carpels and other FM-originating floral organs. In *Galax,* ovules arise from separate ovule primordia along the central axis within the 3-carpellate ovary (axile placentation; Palser 1963), and ovule primordium formation is limited by spacing along that placental axis. Thus, ovule number is limited by ovary length, not by ovary width or the phyllotactic pattern at the FM (Yu et al. 2022).

Though 4x pistils were similarly elongated as other floral organs, ovaries were proportionally shorter and styles proportionally longer. Such minimal ovary elongation was apparently enough to accommodate larger, but not more, ovules. After WGD, 4x ovules may have been biased to size reduction, but that seems unlikely since the coefficient of variation for 2x ovule volumes was identical to that of 4x cytotypes (9.86% and 9.84%, respectively). Larger ovule sizes might increase egg/embryo/seed fitness, but the only way to increase egg number is to increase ovule number, which depends on increasing ovary height (Yu et al. 2020; 2022).

Ovary height in *Galax* is partially constrained by its effect on anther-stigma positioning, since a taller ovary will move the stigma closer to the anther, increasing the potential for self-pollination, unless compensated by changes in style or stamen filament length. 4x filaments were the least elongated and styles the most elongated of all floral organs. In other mixed ploidy species with many ovules (i.e. in species that have not been strongly canalized to produce only one or two ovules), ovule numbers of natural intraspecific autotetraploids relative to diploid progenitors were either the same (Fukuhara 2000; Chansler et al. 2016; Oliveira et al. 2022; García-Muñoz et al. 2023), or slightly fewer (Husband et al. 2002).

### Pollen number increased, but not as a direct effect of WGD

Total allocation to volumes of primary sexual organs (ovules and anthers) in autotetraploid plants was about double (213%) that in diploids. But allocation to anther volume rose from 40% of total primary sex allocation in diploids to 52% in autotetraploids. Altered sex allocation in autotetraploids, suggests that some portion of the anther size increase was compensated by a smaller increase in allocation to ovule sizes, as assumed by sex allocation theory when resources are limiting. A partial physiological trade-off in sex allocation is supported by the fact that 4x anthers had the highest proportional increase in length (39%) relative to other organs, whereas 4x ovaries had near the lowest increase (28%). Furthermore, the total volume of pollen per inflorescence increased far more than the total volume of ovules (147% versus 70%, respectively). Yet, the more-than-doubled allocation to reproduction in autotetraploids relative to diploids suggests they were either not resource-limited or they evolved greater resource acquisition ability, or both. The lack of size-number trade-offs within cytotypes in floral traits suggests resource-limitation was not the primary factor enforcing an apparent trade-off in pre-zygotic sex allocation with the shift to autotetraploidy.

Altered sex allocation can also evolve as a byproduct of differential selection on gamete production when there are evolutionary developmental biases on gamete-bearing organ sizes. Pollen production is linked to anther size, via the size and number of their sporogenous cell precursors, formed in early development of the inner anther (Marchant and Walbot 2022). But WGD increases the size, not the number of pollen grains. To increase the number of pollen grains, either cell cycles that produce sporogenous cells and their descendants must be accelerated or anther maturation must be delayed so that sporogenous cell lineages have more time to undergo more cell divisions. The latter is supported by the fact that increases in genome size prolong the cell cycle (Bennett 1971; Evans et al. 1972; Francis et al. 2008), resulting in phenomena such as delayed flower opening and longer life cycles in some natural autotetraploids (Levin 1983; Corneillie et al. 2019). Yet, tetraploid species have often evolved accelerated pollen cell cycles (shorter duration and more cell cycles per time) compared to their diploid relatives (Bennett and Smith 1972; Finch and Bennett 1972). This seems to be the case in *Galax,* since anthers were already starting to open just before anthesis in both 2x and 4x plants, and 4x plants flowered at similar or earlier times than 2x plants at similar elevations (Barringer and Galloway 2017). That *Galax* autotetraploids have evolved 85% higher pollen production without apparent change in the duration of development suggests both anther and pollen precursor cell cycles became accelerated.

Cell cycle acceleration might evolve by increased biosynthetic capacity due to doubled gene dosage (Conant et al. 2014; Bohutínská et al. 2021), which can affect the speed of cell wall construction and intracellular processes (Bennett 1971; Sugiyama 2005; Williams and Oliveira 2020). At the same time, cell size reduction causes shorter cell cycles to evolve, and pollen of natural *Galax* autotetraploids was 67% smaller than its presumed doubled size at the time of WGD, as is so widely documented in synthetic autotetraploids (see below). Cell size reduction might occur alongside genome downsizing after WGD, but this was not the case in *Galax,* since its 4x genome size was consistently at least double the size of the 2x genome.

In sum, our results suggest the following scenario. Anther enlargement and doubled pollen volume evolved as known effects of WGD. Soon afterwards, both acceleration of sporogenous cell cycles and decreases in cell sizes evolved in concert. After WGD, conserved 4x anther size, not resource limitation, constrains pollen size-number evolution to a near perfect trade-off with a slope near negative one (Figure 4). Enlarged 4x anthers might only appear to be conserved because they evolve slowly relative to pollen number and size, or 4x anther size may be maintained for its capacity to support greater pollen production.

### Polyploidy, breeding systems and pollination efficiency

Our finding that *Galax* diploids had an evolved pollen-ovule ratio of 832 suggests that they are unlikely to be either obligately selfing or obligately outcrossing and is consistent with species that have mixed mating systems (Harder and Johnson 2023). East (1940) reported self-fertility in *Galax,* whereas pollination experiments of Barringer and Galloway (2017) found self-sterility in both cytotypes and genetic patterns indicate low levels of inbreeding (Servick et al. 2015). Retention of self-compatibility in an outcrossing species might explain one puzzling aspect of our findings – 2x stigma-anther distance was conserved in 4x plants, yet it should have increased due to the generally isometric scaling among all other floral organs within and between cytotypes. Both cytotypes seem to have incomplete dichogamy and the capacity for autonomous (within flower) self-pollination, which suggests reproductive assurance by (rare) self-pollination has been advantageous.

Pollen-ovule ratios may be imperfect indicators of breeding systems and the long-term efficiency of pollination (Cruden 1977; Charnov 1982; Queller 1984; Bochynek and Burd 2024), but they can be more meaningfully interpreted when changes in the underlying magnitudes of pollen and ovule numbers are considered (Pellmyr et al. 2020; Cunha et al. 2022; Harder and Johnson 2023). In a progenitor-descendant system such as in *Galax*, autotetraploid populations have evolved a 78% higher PO ratio quite rapidly and entirely as a consequence of changes in pollen number. Both the conservation of ovule number and the increase in pollen number must be explained as post-WGD phenomena, since there is no evidence that WGD itself can cause large changes in cell or organ number. Ovule number may be constrained by development to evolve slowly, as discussed above. Increased pollen production is usually seen as evolving in response to insufficient outcross pollination (Cruden 1977; Aizen and Harder 2007; Alonso et al. 2012) or to increased dispersal distances of pollen relative to seed (Fromhage and Kokko 2010). *Galax* 2x and 4x populations share the same potential pollinators (visitors) and flower concurrently at the same elevation, but seed set in both is outcross pollen limited (Barringer and Galloway 2017). It is also possible that pollen number evolves after WGD by developmental bias during anther development (see next section).

In plants with large bisexual flowers, allocation to secondary sexual organs is generally male-biased, as predicted by sex allocation theory (Paterno et al. 2020; Cunha and Aizen 2023). Similarly, within each *Galax* cytotype, petal length increased with flower (pedicel) size (*β* = 0.30), whereas sepal length did not (*β* = 0). But flower enlargement by WGD is more costly, since WGD did not disproportionally increase petal size relative to sepal size. Male-bias evolves if greater pollinator attraction results in higher fitness returns from increased pollen export than from increased pollen reception. Importantly, we do not know if the larger flowers and displays of *Galax* resulted in greater pollinator attraction or higher male fitness (see Schmickl et al. 2024). Thus, enlarged flower size and presumably pollinator attraction is a direct outcome of WGD, whereas increased pollen production evolves after WGD for other reasons.

### Do polyploids commonly have higher pollen production and if so, why?

Two universal consequences of WGD are the roughly doubled volume of pollen at inception, and the pollen size reduction that follows after WGD (Muller 1979; Butterfass 1987; Williams, unpublished). For example, 2x pollen of synthetic autotetraploids of *Arabidopsis arenosa* was 98% larger in volume than in their diploid parents, but pollen of wild autotetraploids was on average only 9% larger (Westermann et al. 2024). If pollen size and number are developmentally linked through anther size, at least in young polyploids, then the pervasive effect of pollen size reduction after WGD should also cause widespread shifts to higher pollen production as polyploids age. In newly formed synthetic autotetraploids relative to diploid parents, pollen volume was 77-95% larger, but pollen production was 6-32% lower (Baldwin and Husband 2011; Ku et al. 2022). Pollen production in older established cultivars of synthetic autotetraploids ranged from 34% lower to 27% higher than in diploid progenitors (Scott and Longden 1970; Mackiewicz 1965; Sree Rangasamy and Raman 1973). Though sparse, these data support the null hypothesis that WGD itself negatively biases pollen production if at all, since rapid enlargement of pollen size should be compensated by fewer, not greater, numbers of pollen grains. Higher pollen production must evolve soon or long after WGD.

In natural intraspecific autotetraploid populations, pollen number per flower of 4x relative to 2x populations ranged from -8% in *Erysimum* (Garcia-Munoz et al 2023), to -3% in *Corydalis* (Fukuhara 2000), to +5% in *Libidibia* (Oliveira et al. 2022), to +14% in *Chamerion* (Baldwin and Husband 2011), and up to +85% in *Galax* (this study). Diploid pollen grains in all these populations are inferred to have evolved to smaller sizes, losing from 39-68% of their WGD-associated volumes. Thus, in young autotetraploids, size reduction of pollen is not always accompanied by increased pollen production.

A literature review shows similar variation in outcomes for tetraploid species. We compared pollen production among 44 nearest-relative pairs of diploid and present-day tetraploid species in 34 genera (ploidy levels from Rice et al. 2019; pollen production from Cunha et al. 2022) (Supporting Information Table S3). Tetraploids produced fewer pollen grains per flower than diploids in 24 pairs and more pollen in 20 pairs (one-tailed binomial test; *P* = 0.56). The wide range of outcomes in both young, natural autotetraploids and established tetraploid species likely points to a wide range of evolutionary pressures that affect pollen number during polyploid establishment and subsequent evolution, not to a common developmental bias. Polyploids often face outcross pollen limitation during establishment, and WGD provides a novel mechanism for rapidly increasing male fitness by simultaneously increasing pollinator attraction and enabling the post-WGD evolution of higher pollen production without additional post-WGD costs.

## Acknowledgements

The authors thank Highlands Biological Station for lab space and a William Chambers Coker Fellowship in Botanical Research, the UTK herbarium for a Hochmann award, and the UTK Office of Undergraduate Research summer stipends to AK, TB, and CB. We also thank the US National Park Service and the NC Division of Parks and Recreation for granting access.

## Author contributions

JW designed the study; collected, analyzed and interpreted data, and wrote the manuscript. AK, CB, BC, and TB collected data and assisted with interpretation and writing.

## Data availability

Raw data will be deposited on Dryad, https://datadryad.org/

