## Supplemental Figure S1 and Tables S1-3 for "Whole genome duplication enables rapid evolution of male-biased sex allocation in *Galax urceolata*"

**Supplementary Figure 1.** Relative florescence intensity of leaves from putative diploids (2x) and autotetraploids (4x) of *Galax urceolata*. In each panel, leaves of three individuals per putative cytotype were run with a leaf of the standard, *Pisum sativum* var. Ctirad. Note that the 4x peak in A of about 8,000 units is almost exactly double that of the 2x peak of about 4,000 units in C. In B, two individuals from a 2x population and one individual from a 4x population were run together.

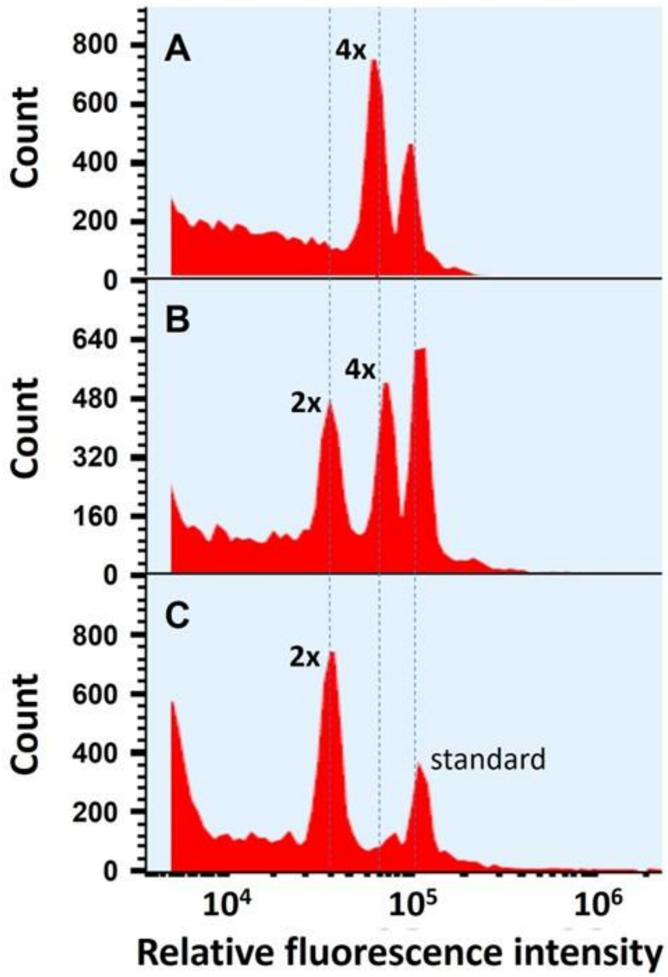

**Supplementary Table 1.** Two-way ANOVA results for whole plant primary sexual allocation to volume in *Galax urceolata*. Organ gender is total anther volume versus ovule volume ($mm^3$). Means and 95% confidence intervals in brackets are back-transformed values.

A) All plants ($N$ = 156), $R^2_{adj.}$ = 0.38.

| Source | DF | Sum of Squares | F Ratio | *P* | Ploidy | Sporangia | Least Sq Mean | Lower 95% | Upper 95% |
|---|---|---|---|---|---|---|---|---|---|
| Cytotype | 1 | 8.7114735 | 170.9524 | **<.0001** | 2x | Ovules | 14.97 | 13.28 | 16.87 |
| Gender | 1 | 0.3207017 | 6.2934 | **0.0126** | 2x | Anthers | 10.13 | 8.99 | 11.42 |
| Cytotype*Gender | 1 | 0.8542193 | 16.763 | **<.0001** | 4x | Ovules | 25.44 | 22.72 | 28.48 |
| | | | | | 4x | Anthers | 27.94 | 24.97 | 31.25 |

B) Plants with one inflorescence only ($N$ = 125), $R^2_{adj.}$ = 0.41.

| Source | DF | Sum of Squares | F Ratio | Prob > F | Ploidy | Sporangia | Least Sq Mean | Lower 95% | Upper 95% |
|---|---|---|---|---|---|---|---|---|---|
| Cytotype | 1 | 5.791851 | 107.5901 | **<.0001** | 2x | Ovules | 13.18 | 11.28 | 15.40 |
| Gender | 1 | 0.084922 | 1.5775 | 0.211 | 2x | Anthers | 10.17 | 8.700 | 11.88 |
| Cytotype*Gender | 1 | 0.17411 | 3.2343 | 0.074 | 4x | Ovules | 27.34 | 22.81 | 32.76 |
| | | | | | 4x | Anthers | 28.63 | 23.95 | 34.22 |

**Supplementary Table 2.** Two-way ANOVA results for whole plant primary sexual allocation to volume in *Galax urceolata*. Gamete gender is total pollen number versus ovule number. Means and 95% confidence intervals in brackets are back-transformed values.

A) All plants ($N = 156$), $R^2_{adj.} = 0.98$.

| Source | DF | Sum of Squares | F Ratio | *P* | Ploidy | Gamete numbers | Least Sq Mean | Lower 95% | Upper 95% |
|---|---|---|---|---|---|---|---|---|---|
| Cytotype | 1 | 1.69813 | 37.6571 | **<.0001** | 2x | Ovules | 3.68 | 3.63 | 3.72 |
| Gender | 1 | 721.1025 | 15990.85 | **<.0001** | 2x | Pollen | 6.60 | 6.55 | 6.65 |
| Cytotype*Gender | 1 | 1.21134 | 26.8621 | **<.0001** | 4x | Ovules | 3.70 | 3.65 | 3.74 |
| | | | | | 4x | Pollen | 6.87 | 6.82 | 6.92 |

B) Plants with one inflorescence only ($N = 125$), $R^2_{adj.} = 0.98$.

| Source | DF | Sum of Squares | F Ratio | Prob > F | Ploidy | Gamete numbers | Least Sq Mean | Lower 95% | Upper 95% |
|---|---|---|---|---|---|---|---|---|---|
| Cytotype | 1 | 1.94316 | 41.9166 | **<.0001** | 2x | Ovules | 3.66 | 3.60 | 3.71 |
| Gender | 1 | 563.6716 | 12159.14 | **<.0001** | 2x | Pollen | 6.55 | 6.50 | 6.61 |
| Cytotype*Gender | 1 | 1.21605 | 26.2318 | **<.0001** | 4x | Ovules | 3.69 | 3.64 | 3.74 |
| | | | | | 4x | Pollen | 6.87 | 6.82 | 6.92 |

**Supplementary Table 3.** Comparison of pollen numbers per flower for nearest-relative clades. Values are $\log_{10} \pm 1 \log_{10}$ standard deviation. Where there was data for more than one species in a sister clade (2x or 4x), values were averaged. 1x is the haploid number of chromosomes in the 2x taxon, which is the base chromosome number for that genus (all 4x taxa had twice as many chromosomes). Effect size is $\log_{10}$(4x) minus $\log_{10}$(2x) pollen number.

| Genus | 1x | 2x clade | $\log_{10}$ 2x pollen no. | 4x clade | $\log_{10}$ 4x pollen no. | $\log_{10}$ effect size |
|---|---|---|---|---|---|---|
| *Allium* | 8 | *Allium wallichii* | 4.62 | *Allium cyaneum* | 4.54 | -0.08 |
| *Alyssum* | 8 | *Alyssum linifolium, A. loiseleurii, A.simplex* | 3.36 ± 0.74 | *Alyssum damscenum* | 2.79 | -0.57 |
| *Anagallis* | 10 | *Anagallis monelli* | 4.76 | *Anagallis arvensis* | 4.20 | -0.56 |
| *Anemone* | 8 | *Anemone cylindrica* | 4.98 | *Anemone multifida* | 4.05 | -0.93 |
| *Arabis* | 8 | *Arabis aucheri* | 2.99 | *Arabis verna* | 3.27 | 0.29 |
| *Arum* | 14 | *Arum cylindraceum* | 3.83 | *Arum maculatum* | 3.74 | -0.10 |
| *Callistemon* | 11 | *Callistemon citrinus* | 4.29 | *Callistemon viminalis* | 5.11 | 0.82 |
| *Capsella* | 8 | *Capsella grandiflora, C. rubella* | 4.20 ± 0.35 | *Capsella bursa-pastoris* | 3.91 | -0.29 |
| *Conophytum* | 9 | *Conophytum luckhoffii, C. pageae* | 5.33 ± 0.43 | *Conophytum flavum* | 5.35 | 0.02 |
| *Crotalaria* | 8 | *Crotalaria micans* | 5.22 | *Crotalaria pumila, C. stipularia* | 5.10 ± 0.44 | -0.12 |
| *Delphinium* | 8 | *Delphinium emarginatum, D. fissum, D. sylvaticum* | 5.14 ± 0.15 | *Delphinium montanum* | 4.94 | -0.21 |
| *Dianthus* | 15 | *Dianthus superbus* | 4.50 | *Dianthus hyssopifolius* | 4.44 | -0.05 |
| *Erodium n=9* | 9 | *Erodium ciconium* | 2.65 | *Erodium chrysanthum* | 3.29 | 0.64 |
| *Erodium n=10* | 10 | *Erodium recoderi, E. rupicola* | 3.73 ± 0.04 | *Erodium acaule, E. cicutarium* | 3.09 ± 0.71 | -0.64 |
| *Erodium n=10* | 10 | *Erodium neuradifolium* | 3.46 | *Erodium nervulosum* | 2.72 | -0.74 |
| *Erodium n=10* | 10 | *Erodium chium* | 3.19 ± 2.93 | *Erodium malacoides* | 2.93 | -0.26 |
| *Erodium n=10* | 10 | *Erodium carvifolium* | 3.58 | *Erodium manescavi* | 3.55 | -0.03 |
| *Erythronium* | 12 | *Erythronium grandiflorum* | 4.91 | *Erythronium americanum* | 5.00 | 0.09 |
| *Genista* | 12 | *Genista cinerascens, G. polyanthos* | 4.16 ± 0.04 | *Genista florida* | 4.29 | 0.13 |
| *Helianthemum* | 5 | *Helianthemum squamatum* | 3.58 | *Helianthemum ovatum* | 4.42 | 0.85 |
| *Indigofera* | 8 | *Indigofera suffruticosa* | 4.71 | *Indigofera parodiana* | 4.44 | -0.28 |

Table S3 continued…

| | | | | | | |
|---|---|---|---|---|---|---|
| *Litsea* | 12 | *Litsea cubeba* | 4.21 | *Litsea glutinosa* | 3.97 | -0.24 |
| *Mazus* | 10 | *Mazus miquelii, M. stachydifolius* | 4.28 $\pm$ 0.05 | *Mazus pumilus* | 3.91 | -0.37 |
| *Medicago* | 8 | *Medicago edgeworthii* | 3.25 | *Medicago arborea* | 3.73 | 0.48 |
| *Mollugo* | 9 | *Mollugo cerviana* | 2.9 | *Mollugo pentaphylla* | 2.98 | 0.08 |
| *Opuntia* | 11 | *Opuntia microdasys* | 4.75 | *Opuntia robusta* | 5.41 | 0.67 |
| *Passiflora* | 6 | *Passiflora misera* | 4.38 | *Passiflora suberosa* | 3.8 | -0.58 |
| *Phyllanthus* | 13 | *Phyllanthus niruri* | 2.76 | *Phyllanthus urinaria* | 2.47 | -0.29 |
| *Pinguicula* | 8 | *Pinguicula villosa* | 3.86 | *Pinguicula alpina* | 5.08 | 1.22 |
| *Scrophularia* | 13 | *Scrophularia canina* | 5.43 | *Scrophularia herminii* | 5.88 | 0.45 |
| *Securigera* | 6 | *Securigera securidaca* | 4.28 | *Securigera varia* | 4.37 | 0.09 |
| *Senecio* | 20 | *Senecio portalesianus* | 3.41 | *Senecio bipontinii* | 3.57 | 0.15 |
| *Silene* | 12 | *Silene viscosa* | 4.66 | *Silene caroliniana, S. stellata, S. virginica* | 4.42 $\pm$ 0.28 | -0.24 |
| *Silene* | 12 | *Silene spergulifolia* | 4.38 | *Silene vallesia* | 4.48 | 0.10 |
| *Suaeda* | 9 | *Suaeda monoica* | 4.21 | *Suaeda maritima* | 4.28 | 0.07 |
| *Tillandsia* | 25 | *Tillandsia recurvata* | 3.23 | *Tillandsia tricholepis* | 3.09 | -0.13 |
| *Trifolium* | 8 | *Trifolium alpinum* | 3.99 | *Trifolium repens* | 3.63 | -0.36 |
| *Vaccinium* | 12 | *Vaccinium vitis-idaea* | 4.56 | *Vaccinium oxycoccos* | 4.7 | 0.14 |
| *Veronica n=7* | 7 | *Veronica syriaca* | 4.05 | *Veronica bozakmanii* | 2.72 | -1.33 |
| *Veronica n=7* | 7 | *Veronica polita* | 3.61 | *Veronica persica* | 3.64 | 0.03 |
| *Veronica n=8* | 8 | *Veronica arvensis* | 2.72 | *Veronica chamaedrys* | 4.08 | 1.37 |
| *Veronica n=9* | 9 | *Veronica cusickii* | 4.00 | *Veronica bellidioides* | 3.96 | -0.03 |
| *Veronica n=9* | 9 | *Veronica lycica* | 4.01 | *Veronica cymbalaria* | 3.03 | -0.98 |
| *Veronica n=9* | 9 | *Veronica scardica* | 3.44 | *Veronica anagallis-aquatica* | 3.59 | 0.15 |

*Notes:* Nearest-relative diploid-tetraploid relationships are based on genus-level phylogenies used in Rice et al. (2019), except for *Conophytum* (Powell et al. 2022). Autotetraploids and allotetraploids were not distinguished. $Log_{10}$ effect sizes were normally distributed with a mean of -0.035 with a 95% confidence interval that includes zero {-0.199, 0.129}. Chromosome numbers were checked on the Chromosome counts database (Rice et al. 2015).
